# Neuropeptide T2A-GAL4 knock-ins illustrate practical considerations for interpreting GAL4 reporter patterns in *Drosophila*

**DOI:** 10.64898/2026.09.16.750837

**Authors:** Kyoko Jinnai, Kota Ozawa, Katsuki Yajima, Hiromu Tanimoto, Shu Kondo

**Affiliations:** Department of Biological Science and Technology, Tokyo University of Science, Tokyo, Japan; Graduate School of Life Sciences, Tohoku University, Sendai, Japan; Department of Biology, Faculty of Science, Tohoku University, Sendai, Japan

**Keywords:** neuropeptides, *Drosophila*, genome editing, GAL4

## Abstract

Transgenic GAL4 lines are widely used in *Drosophila* to label and manipulate specific cell types, but the resulting reporter expression patterns do not always represent ongoing driver activity. We generated a collection of neuropeptide T2A-GAL4 knock-ins and characterized their expression by native-fluorescence imaging using *UAS-CD8::GFP* and a *brp-miRFP680* knock-in, eliminating the need for immunostaining. During characterization, we observed both timing-dependent changes in adult GFP labeling and GAL4-associated cellular abnormalities. Using temperature-sensitive GAL80, we found that adult GFP labeling in certain neurons driven by *CCHa2-T2A-GAL4* and *Dsk-T2A-GAL4* depended on GAL4 activity before eclosion and was not reproduced by adult-restricted induction. Several GAL4 lines also showed dominant developmental phenotypes. Heterozygous *Burs-T2A-GAL4* animals, for example, displayed wing expansion defects that were fully suppressed by GAL80. Severe wing defects were not accompanied by a reduction in the number of reporter-labeled neurons before eclosion, but the morphology of these neurons was affected. These observations illustrate how developmental driver history and GAL4-associated cellular effects can influence reporter patterns in the adult nervous system.

## 1 Introduction

Neuropeptides regulate diverse aspects of development, physiology and behavior in *Drosophila*, often through small and anatomically defined populations of neurons and endocrine cells (Nässel and Winther 2010; Nässel and Zandawala 2019). Genetic access to these cells is therefore important for dissecting the functions of individual neuropeptides. The GAL4/UAS system has been widely used for this purpose, allowing specific cells to be visualized and manipulated in vivo (Brand and Perrimon 1993). Despite its versatility, however, several features of the GAL4/UAS system can complicate the interpretation of reporter expression patterns and associated phenotypes. First, when expressed at high levels, GAL4 itself can cause developmental defects, apoptosis, neuronal dysfunction and other physiological abnormalities (Kramer and Staveley 2003; Rezával et al. 2007; Keith et al. 2026). Second, GAL4-driven fluorescent reporters do not necessarily provide a faithful readout of ongoing GAL4 activity because stable reporter proteins can persist after GAL4 activity has declined (Manning et al. 2012; Soustelle and Giangrande 2007). Although both issues are well recognized, their practical impact on anatomical reporter patterns remains incompletely characterized, particularly in post-mitotic neurons.

A separate factor affecting the interpretation of GAL4 reporter patterns is the design of the driver itself. Promoter-GAL4 fusions have been generated for many neuropeptide genes (Renn et al. 1999; Park et al. 2008), but such transgenes may lack regulatory elements required to fully reproduce endogenous expression patterns. By contrast, endogenous T2A-GAL4 knock-ins are expected to more closely recapitulate endogenous expression patterns because GAL4 is expressed within the native regulatory context of the targeted gene (Diao and White 2012; Kondo et al. 2020). We therefore extended our previous T2A-GAL4 knock-in strategy to a set of neuropeptide genes in *Drosophila*.

Here, we generated a collection of neuropeptide T2A-GAL4 knock-in lines and characterized their expression in the *Drosophila* nervous system. To facilitate rapid whole-brain imaging, we also generated fluorescent protein knock-ins of the synaptic protein Bruchpilot (Wagh et al. 2006), enabling neuropil landmarks and GAL4-dependent reporters to be visualized entirely by native fluorescence. During characterization of these resources, we encountered several unexpected limitations of GAL4-based labeling. Strong GAL4 activity produced developmental and cellular abnormalities that were suppressible by GAL80 but were not fully explained by p35-sensitive apoptosis. Temporal restriction of GAL4 activity further revealed that transient GAL4 activity in specific cells during the pupal stage produced GFP labeling that remained detectable in the adult brain for at least 5 days. Thus, this study provides useful genetic and imaging resources while also revealing factors that can complicate the interpretation of GAL4-dependent reporter patterns in the adult nervous system.

## 2 Results

### 2.1 Generation of neuropeptide-GAL4 knock-in lines and fluorescent Brp reference alleles

As part of our ongoing effort to generate GAL4 knock-in lines for genes involved in neurotransmission (Kondo et al. 2020), we targeted a set of 21 neuropeptide loci in the *Drosophila* genome (Table 1). For each locus, a T2A-GAL4 cassette was inserted immediately upstream of the endogenous stop codon by CRISPR/Cas9-mediated targeted integration, placing GAL4 in frame with the neuropeptide precursor coding sequence (Figure 1H). In this configuration, GAL4 is transcribed as part of the endogenous transcript, whereas T2A-mediated ribosome skipping allows the neuropeptide precursor and GAL4 to be translated as separate polypeptides (Diao and White 2012; Donnelly et al. 2001). Unless otherwise indicated, the *3xP3-RFP* marker cassette was subsequently removed by Cre-mediated excision.

**Figure 1.**
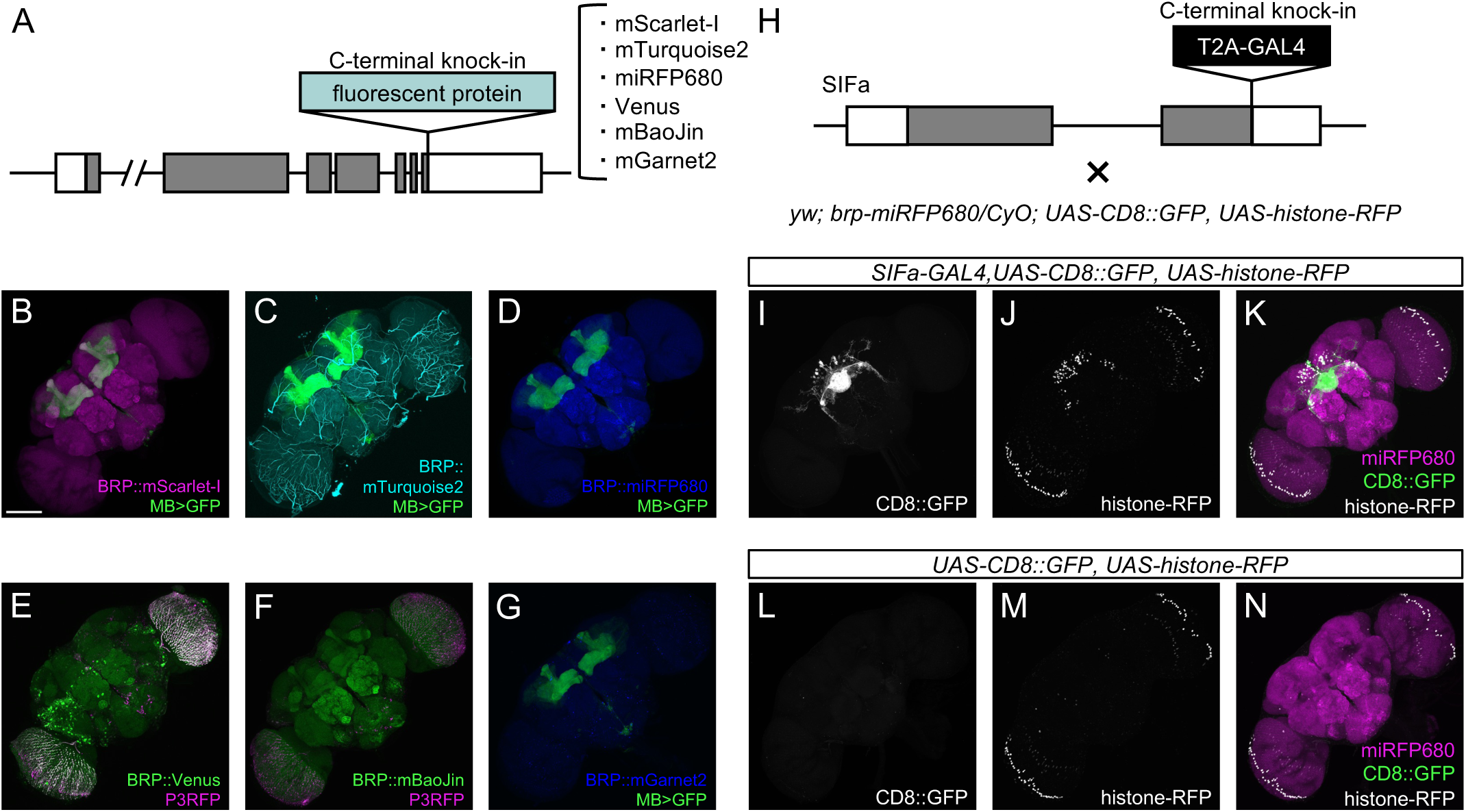
Generation of a multicolor Brp fluorescent knock-in series and neuropeptide-GAL4 knock-in lines. (A) Schematic of the fluorescent knock-in strategy at the endogenous *brp* locus. Fluorescent protein coding sequences were inserted immediately upstream of the *brp* stop codon to generate C-terminal Brp fluorescent fusion alleles. (B–D, G) Representative adult brain images showing R13F02-GAL4-driven CD8::GFP labeling together with Brp-mScarlet-I (B), Brp-mTurquoise2 (C), Brp-miRFP680 (D), or Brp-mGarnet2 (G). (E,F) Fluorescence from Brp-Venus (E) and Brp-mBaoJin (F) was imaged together with RFP from the retained *3xP3-RFP* marker. (H) Schematic of the neuropeptide-GAL4 knock-in strategy. A T2A-GAL4 cassette was inserted immediately upstream of the stop codon of the endogenous neuropeptide coding sequence. The *SIFa* locus is shown as an example. (I–K) Representative adult brain images of *SIFa-T2A-GAL4* reporter labeling using *UAS-CD8::GFP* and *UAS-histone-RFP*. Panels show (I) CD8::GFP-labeled neuronal membranes, (J) histone-RFP-labeled nuclei, and (K) both reporter signals together with Brp-miRFP680-labeled neuropil. (L–N) Corresponding no-GAL4 controls showing (L) CD8::GFP, (M) histone-RFP, and (N) both reporter channels together with Brp-miRFP680-labeled neuropil. Adult brains were obtained from 5-day-old female flies. Scale bar, 100 μm.

**Table 1.** Neuropeptide T2A-GAL4 knock-in lines generated in this study.

| Gene symbol [targeted transcript] | Gene name | CG number |
| --- | --- | --- |
| *Akh* | Adipokinetic hormone | CG1171 |
| *Burs* | Bursicon | CG13419 |
| *CCHa1* | CCHamide-1 | CG14358 |
| *CCHa2* | CCHamide-2 | CG14375 |
| *Crz* | Corazonin | CG3302 |
| *Dh44* | Diuretic hormone 44 | CG8348 |
| *Ms (Dms)* | Myosuppressin | CG6440 |
| *Dsk* | Drosulfakinin | CG18090 |
| *ETH* | Ecdysis triggering hormone | CG18105 |
| *FMRFa* | FMRFamide | CG2346 |
| *Gpb5* | Glycoprotein hormone beta 5 | CG40041 |
| *Hug* | Hugin | CG6371 |
| *ITP [RE]* | Ion transport peptide | CG13586 |
| *Lk* | Leucokinin | CG13480 |
| *NPF* | Neuropeptide F | CG10342 |
| *Nplp4* | Neuropeptide-like precursor 4 | CG15361 |
| *Orcokinin [RA]* | Orcokinin | CG13565 |
| *Pburs* | Partner of Bursicon | CG15284 |
| *Proc* | Proctolin | CG7105 |
| *SIFa* | SIFamide | CG33527 |
| *Trissin* | Trissin | CG14871 |

To facilitate imaging of neuropeptide-GAL4 reporter patterns in the brain, we developed a native-fluorescence strategy that eliminated the need for the standard Brp immunostaining. For whole-brain imaging, the nc82 monoclonal antibody, which recognizes the synaptic protein Bruchpilot (Brp), is widely used to visualize neuropil architecture (Laissue et al. 1999; Wagh et al. 2006); however, the staining procedure typically requires several days of antibody incubation and washing (Kohl et al. 2014). We therefore generated C-terminal Brp fluorescent fusions by phiC31-mediated cassette exchange (Kondo et al. 2020) of the existing *brp-GFP11* allele, replacing the GFP11 cassette with fluorescent protein coding sequences immediately upstream of the endogenous *brp* stop codon (Figure 1A). The resulting alleles encoded Brp fused to mTurquoise2 (cyan), mBaoJin (green), Venus (yellow-green), mScarlet-I (red), mGarnet2 (far-red), or miRFP680 (near-infrared) (Goedhart et al. 2012; Zhang et al. 2024; Nagai et al. 2002; Bindels et al. 2017; Matela et al. 2017; Matlashov et al. 2020). All six alleles produced clean neuropil-specific fluorescence in the larval central nervous system (CNS), adult brain and adult ventral nerve cord and enabled multicolor imaging together with GFP- or RFP-based reporters (Figures 1B–G and S1). Among these proteins, mBaoJin is a recently developed green fluorescent protein reported to exhibit high brightness and exceptional photostability (Zhang et al. 2024). Under matched imaging conditions, Brp-mBaoJin was brighter than Brp-Venus and substantially more resistant to photobleaching (Figure S2). For rapid characterization of the neuropeptide-GAL4 collection, we selected Brp-miRFP680 because its far-red fluorescence could be combined with *UAS-CD8::GFP* and *UAS-histone-RFP*. These three signals simultaneously visualized neuropil landmarks, the membranes and processes of GAL4-expressing cells, and their nuclei, respectively, without immunostaining (Figure 1I–N).

We crossed each neuropeptide-GAL4 knock-in line to *UAS-CD8::GFP* and imaged adult brains together with Brp-miRFP680. This generated an adult brain reference image set for the neuropeptide-GAL4 collection, with GAL4-dependent membrane labeling shown relative to Brp-labeled neuropil landmarks (Figure 2). Corresponding reference image sets were also generated for the larval CNS and adult ventral nerve cord (VNC) (Figures S3 and S4).

**Figure 2.**
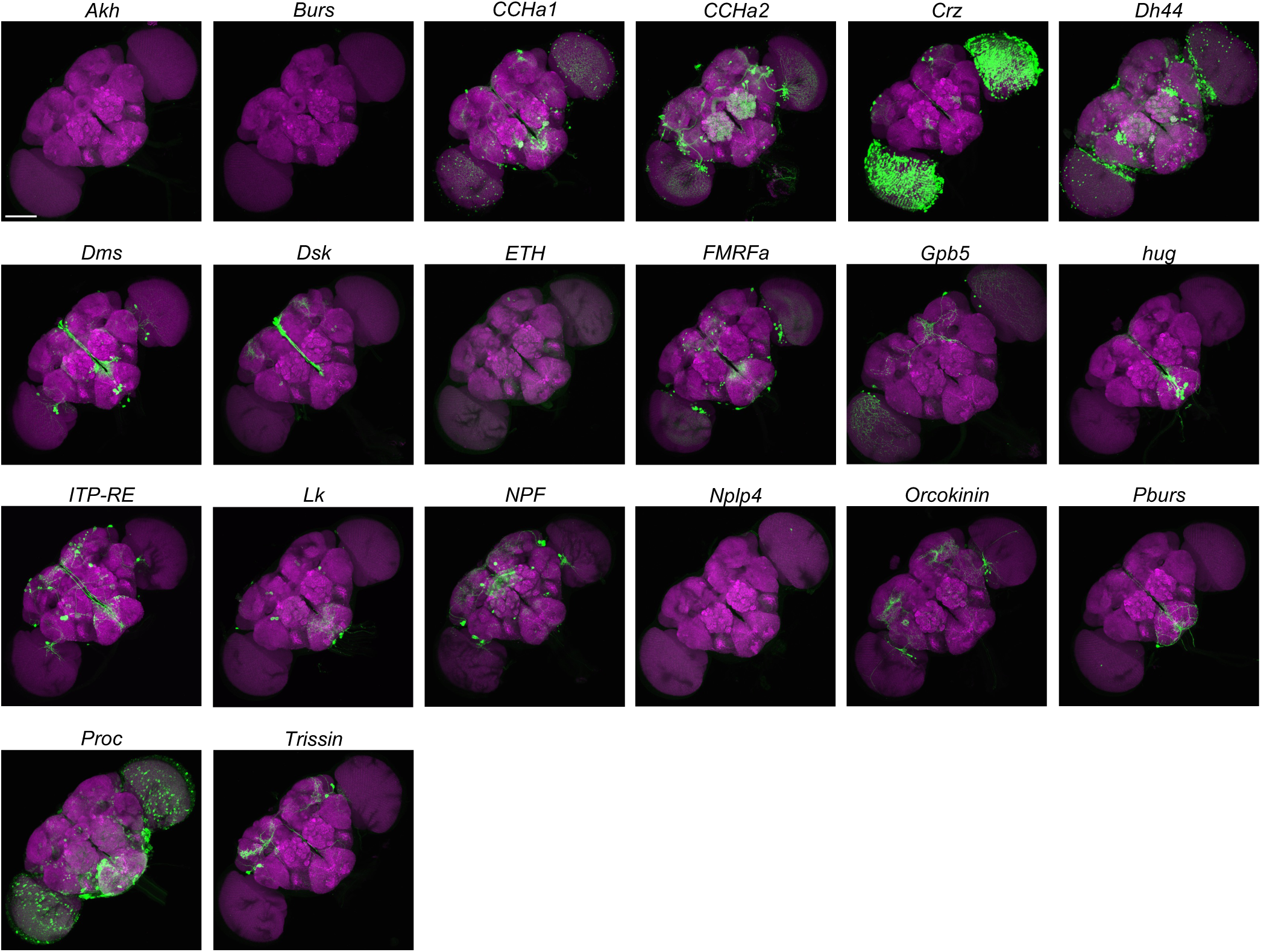
Adult brain reporter labeling patterns of neuropeptide-GAL4 knock-in lines. Representative adult brain montage of the indicated neuropeptide-GAL4 lines. CD8::GFP (green) provides GAL4-dependent membrane labeling, while Brp-miRFP680 (magenta) provides neuropil landmarks. Adult brains were obtained from 5-day-old female flies. Scale bar, 100 μm.

### 2.2 Burs/Pburs knock-in GAL4 lines show GAL80-suppressible wing expansion defects

During the establishment of the neuropeptide-T2A-GAL4 knock-in collection, we noticed prominent wing expansion defects in the *Burs*- and *Pburs*-T2A-GAL4 stocks. The *Bursicon* (*Burs*) and *Partner of Bursicon* (*Pburs*) genes encode the two subunits of Bursicon, a heterodimeric neuropeptide that triggers the expansion and sclerotization of the wing at the end of eclosion. In mutant flies lacking *Burs* or *Pburs*, wings remain folded as in pupae (Dewey et al. 2004; Luo et al. 2005). Interestingly, *Burs*- and *Pburs*-T2A-GAL4 knock-ins dominantly induced the folded-wing phenotype, suggesting that it was not caused by loss of endogenous Bursicon. Thus, we decided to explore the mechanism by which the knock-in alleles affect neuronal function and induce the observed phenotype. We first classified the wing expansion defect into four categories based on severity (Figure 3A).

**Figure 3.**
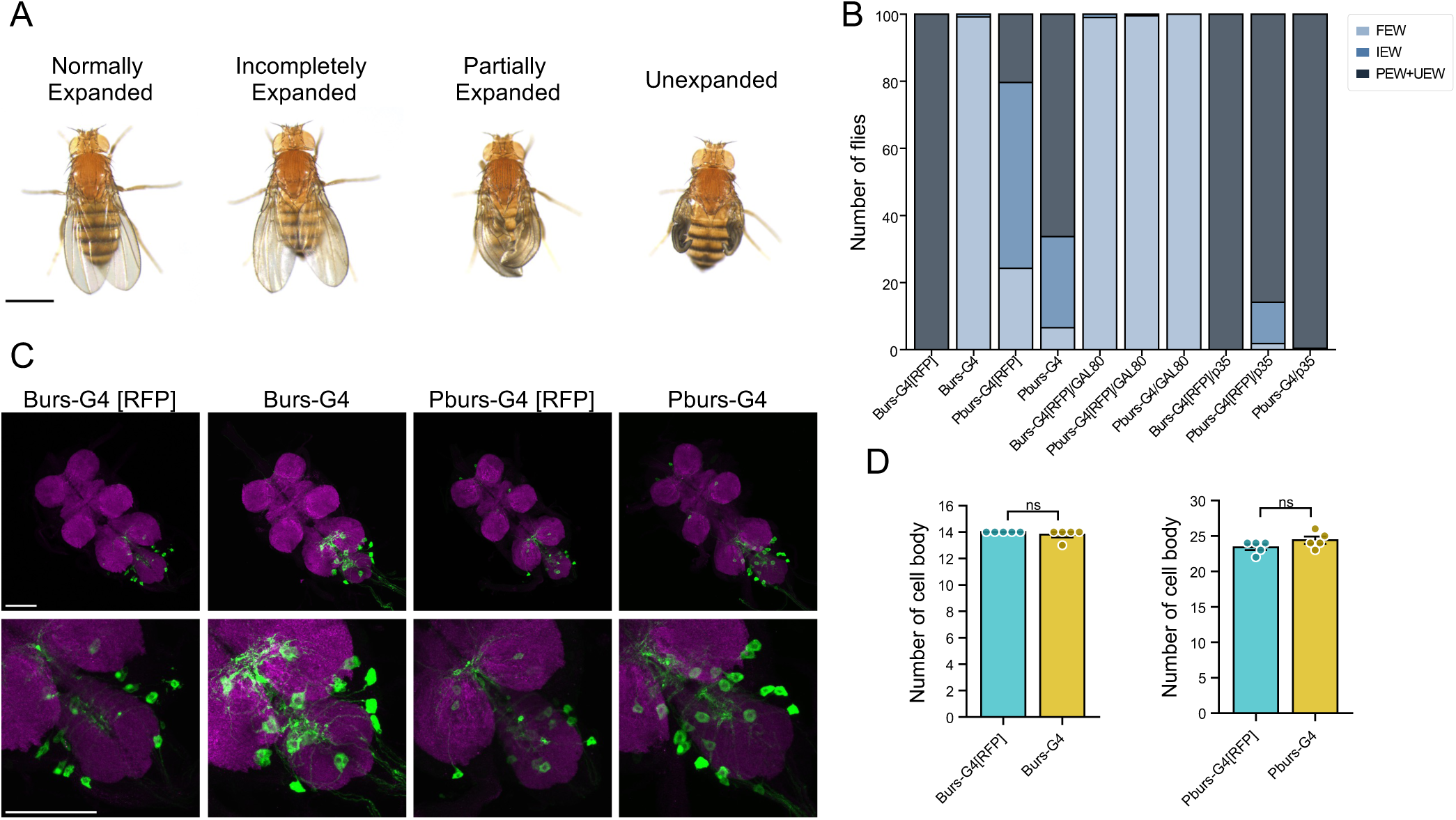
GAL4-dependent wing expansion defects in *Burs/Pburs* knock-in GAL4 lines. (A) Representative adult wing phenotypes used for scoring. Wings were classified as Normally Expanded, Incompletely Expanded, Partially Expanded, or Unexpanded. Scale bar, 1 mm. (B) Distribution of adult wing phenotypes in *Burs-T2A-GAL4* and *Pburs-T2A-GAL4* flies with the indicated genetic backgrounds. Partially Expanded and Unexpanded wings were grouped as severe wing expansion defects. [RFP] indicates alleles retaining the *3xP3-RFP* marker; alleles without [RFP] had the marker removed by Cre-mediated excision. *tubP-GAL80* was used to suppress GAL4 activity, and *UAS-p35* to inhibit p35-sensitive apoptosis. Bars show the distribution of wing phenotype categories (n = 184–466 flies per genotype). Both males and females were scored. (C) Representative pre-eclosion VNC images of *Burs-T2A-GAL4* and *Pburs-T2A-GAL4* flies. CD8::GFP (green) provides GAL4-dependent membrane labeling, while Brp-miRFP680 (magenta) provides neuropil landmarks. Scale bar, 100 μm. (D) Quantification of the number of reporter-labeled cell bodies in individual VNCs of the genotypes shown in (C). Data are presented as mean ± SEM with individual data points; n = 5 VNCs per genotype. Marker-retained and marker-excised alleles were compared separately for each driver using Welch’s *t*-test; ns, not significant.

Using this scoring scheme, we examined wing phenotypes in *Burs*- and *Pburs-T2A-GAL4* animals. Nearly all *Burs-T2A-GAL4[RFP]* flies, where [RFP] denotes retention of the *3xP3-RFP* marker, showed severe wing expansion defects, whereas *Pburs-T2A-GAL4* flies showed milder wing expansion phenotypes regardless of whether the *3xP3-RFP* marker was retained (Figure 3B). These wing expansion defects were completely suppressed by co-expression of the GAL4 inhibitor GAL80 (Lee and Luo 1999), indicating that they depended on GAL4 activity. Interestingly, the wing expansion phenotype of *Burs-T2A-GAL4* animals was largely suppressed by removal of the *3xP3-RFP* selection marker from the knock-in cassette. In the original cassette, GAL4 is followed by a heterologous polyadenylation signal that terminates the transcript upstream of the endogenous 3′ UTR (Kondo et al. 2020). Cre-mediated excision removes this polyadenylation signal together with the *3xP3-RFP* marker, allowing GAL4 transcripts to use the native 3′ UTR and polyadenylation signal of the target gene. This suggests that the heterologous polyadenylation signal in the knock-in cassette enhanced GAL4 activity, possibly through increased transcript stability or translational efficiency.

Because excessive GAL4 activity is known to induce apoptosis (Abidi and Smith-Bolton 2018; Kramer and Staveley 2003; Rezával et al. 2007), we next asked whether the wing expansion defects could be suppressed by the caspase inhibitor p35 (Hay et al. 1994). Expression of *UAS-p35* failed to restore wing expansion in *Burs-T2A-GAL4[RFP]* animals and produced only a slight improvement in *Pburs-T2A-GAL4* animals (Figure 3B). Thus, p35-sensitive cell death is unlikely to account for the GAL4-induced wing expansion defects. Indeed, when *Burs-* or *Pburs-T2A-GAL4* was used to drive *UAS-CD8::GFP*, the positions of GFP-positive VNC neurons in late pupae were consistent with the previously described distribution of Bursicon-expressing neurons (Figure 3C,D; Luan et al. 2006; Peabody et al. 2008). In particular, while removal of the *3xP3-RFP* marker markedly rescued the wing phenotype of *Burs-T2A-GAL4* animals, it did not alter the number of GFP-labeled neurons. However, in the marker-retained line, the cell bodies appeared smaller and the arbors less extensive than in the marker-excised line, suggesting that excessive GAL4 activity may impair neuronal morphology or physiology without simply eliminating the labeled cells.

### 2.3 Apoptosis-independent effects of GAL4 activity in multiple drivers

In addition to the neuropeptide loci, several other T2A-GAL4 knock-ins that we generated exhibited striking dominant phenotypes. During our previous effort to generate T2A-GAL4 knock-ins for neurotransmission-related genes (Kondo et al. 2020), we were unable to establish balanced stocks for *Gad1* and *Tdc2*, although multiple independent transformants were obtained. Heterozygous *Gad1-T2A-GAL4* animals invariably died during pupal development, whereas females heterozygous for *Tdc2-T2A-GAL4* were sterile. Because loss-of-function alleles of these genes do not cause the corresponding phenotypes in heterozygotes (Cole et al. 2005; Featherstone et al. 2000), these dominant effects are consistent with toxicity associated with GAL4 expression rather than disruption of the targeted locus. Notably, homozygous *Tdc2* loss-of-function mutants are female sterile (Cole et al. 2005), supporting the idea that the dominant sterility of *Tdc2-T2A-GAL4* reflects selective impairment of *Tdc2*-expressing cells. Together with the phenotypes observed in the neuropeptide knock-ins, these observations suggest that deleterious effects of GAL4 activity may occur in neuronal drivers more frequently than is generally appreciated.

Adverse effects of GAL4 on development and physiology in *Drosophila* have previously been reported for strong drivers, including *GMR-GAL4* and ubiquitous GAL4 drivers, and apoptosis has often been implicated as an underlying mechanism (Kramer and Staveley 2003). We therefore asked whether apoptosis also accounted for the strong developmental phenotype of *nub-T2A-GAL4*, which we generated alongside the neuropeptide knock-ins. In heterozygous *nub-T2A-GAL4* animals, GAL4 is expressed predominantly in the larval wing imaginal disc (Figure S5; Cifuentes and García-Bellido 1997), and the resulting adult wings were smaller and showed conspicuous defects in the wing margin (Figure 4A). Co-expression of GAL80 restored essentially normal wing morphology, confirming that the phenotype depended on GAL4 activity. In contrast, expression of p35 failed to rescue the wing phenotype and instead exacerbated some aspects of the defect. Consistent with this observation, immunostaining of larval wing imaginal discs for cleaved Dcp-1 did not reveal a marked increase in apoptosis in *nub-T2A-GAL4* compared with control discs (Figure S5). Thus, as in Bursicon-positive neurons, the developmental abnormalities caused by GAL4 could not be explained simply by p35-sensitive apoptosis.

**Figure 4.**
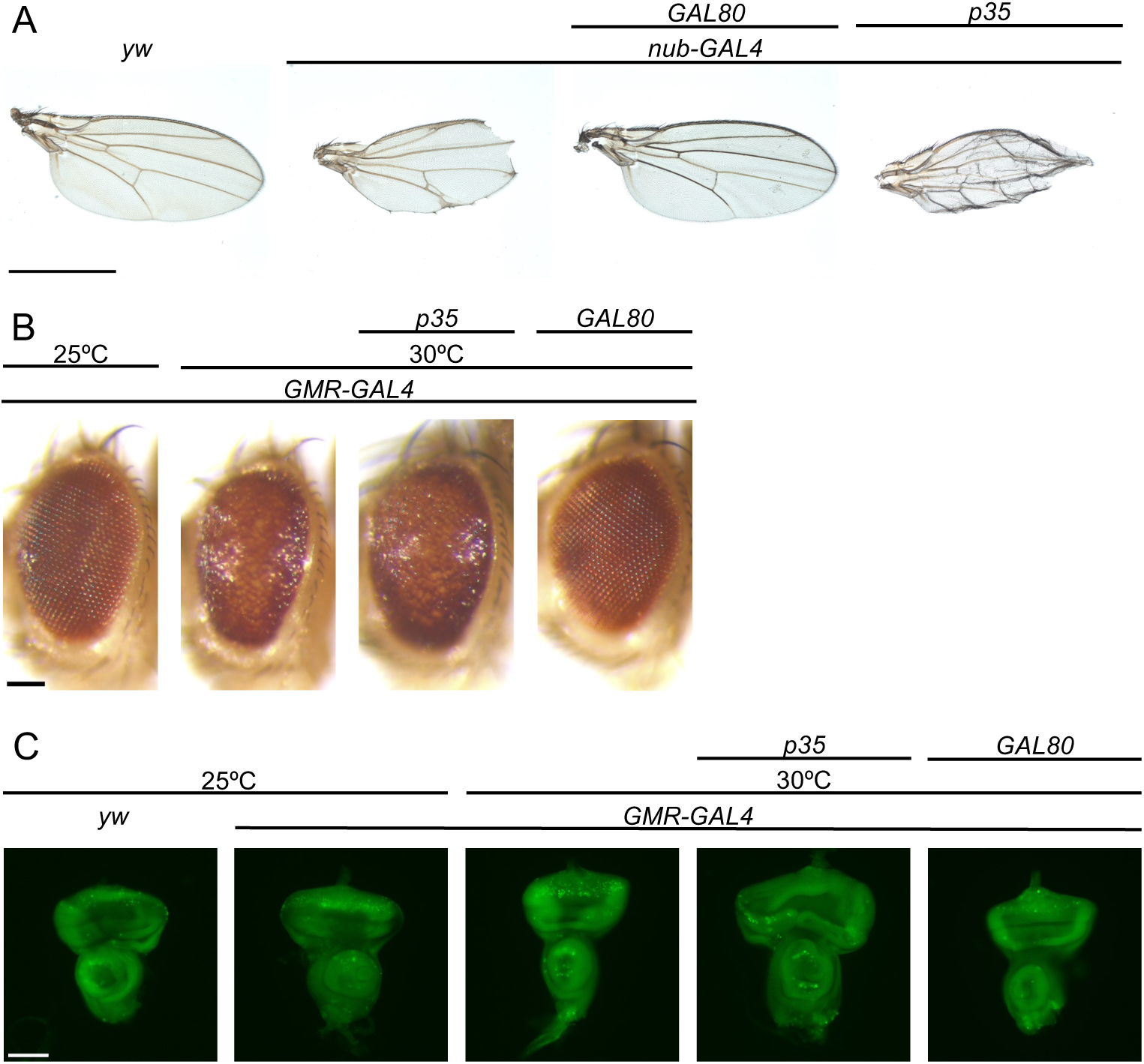
GAL4-dependent morphological defects in *nub-T2A-GAL4* and *GMR-GAL4* flies. (A) Representative adult wings of female *yw*, *nub-T2A-GAL4*, *nub-T2A-GAL4/tubP-GAL80*, and *nub-T2A-GAL4/UAS-p35* flies reared at 25°C. Scale bar, 500 μm. (B) Representative adult eyes of female *GMR-GAL4* flies reared at 25°C, and *GMR-GAL4*, *GMR-GAL4/UAS-p35*, and *GMR-GAL4/tubP-GAL80* flies reared at 30°C. Scale bar, 100 μm. (C) Acridine Orange staining of third-instar larval eye imaginal discs from *yw* and *GMR-GAL4* flies reared at 25°C, and *GMR-GAL4*, *GMR-GAL4/UAS-p35*, and *GMR-GAL4/tubP-GAL80* flies reared at 30°C. Scale bar, 100 μm. *tubP-GAL80* was used to suppress GAL4 activity, and *UAS-p35* was used to inhibit p35-sensitive apoptosis.

We next revisited *GMR-GAL4*, for which GAL4-induced apoptosis in the developing eye has been well documented. At 30°C, a single copy of *GMR-GAL4* induced elevated apoptosis posterior to the morphogenetic furrow and produced a rough-eye phenotype in adults (Figure 4B,C). GAL80 suppressed both apoptosis in the eye disc and the adult rough-eye phenotype. By contrast, p35 almost completely suppressed cell death posterior to the morphogenetic furrow in the eye disc but did not restore adult eye morphology. Thus, although apoptosis is clearly induced by *GMR-GAL4*, suppression of this cell death is insufficient to prevent the final morphological defect. These results indicate that strong GAL4 activity can impair cellular morphology and function through mechanisms other than p35-sensitive apoptosis.

### 2.4 Adult-restricted GAL4 can generate different labeling patterns in the brain

Given the apparent prevalence of GAL4-associated effects on cellular morphology and survival, we next asked whether continuous GAL4 activity altered the presence or morphology of GAL4-expressing cells in any of our neuropeptide knock-in lines. We used temperature-sensitive GAL80 (GAL80ts) to restrict GAL4 activity to a narrow time window in adulthood to minimize the adverse effect of GAL4 (McGuire et al. 2003). Each neuropeptide T2A-GAL4 knock-in line was crossed to a tester strain carrying *tubP-GAL80ts*, *UAS-CD8::GFP*, and *UAS-histone-RFP*. Progeny were reared at 18°C to suppress GAL4 activity during development and then shifted to 30°C 4–6 days after eclosion to activate GAL4. Adult brains were examined 2 days after the temperature shift (Figure 5A).

**Figure 5.**
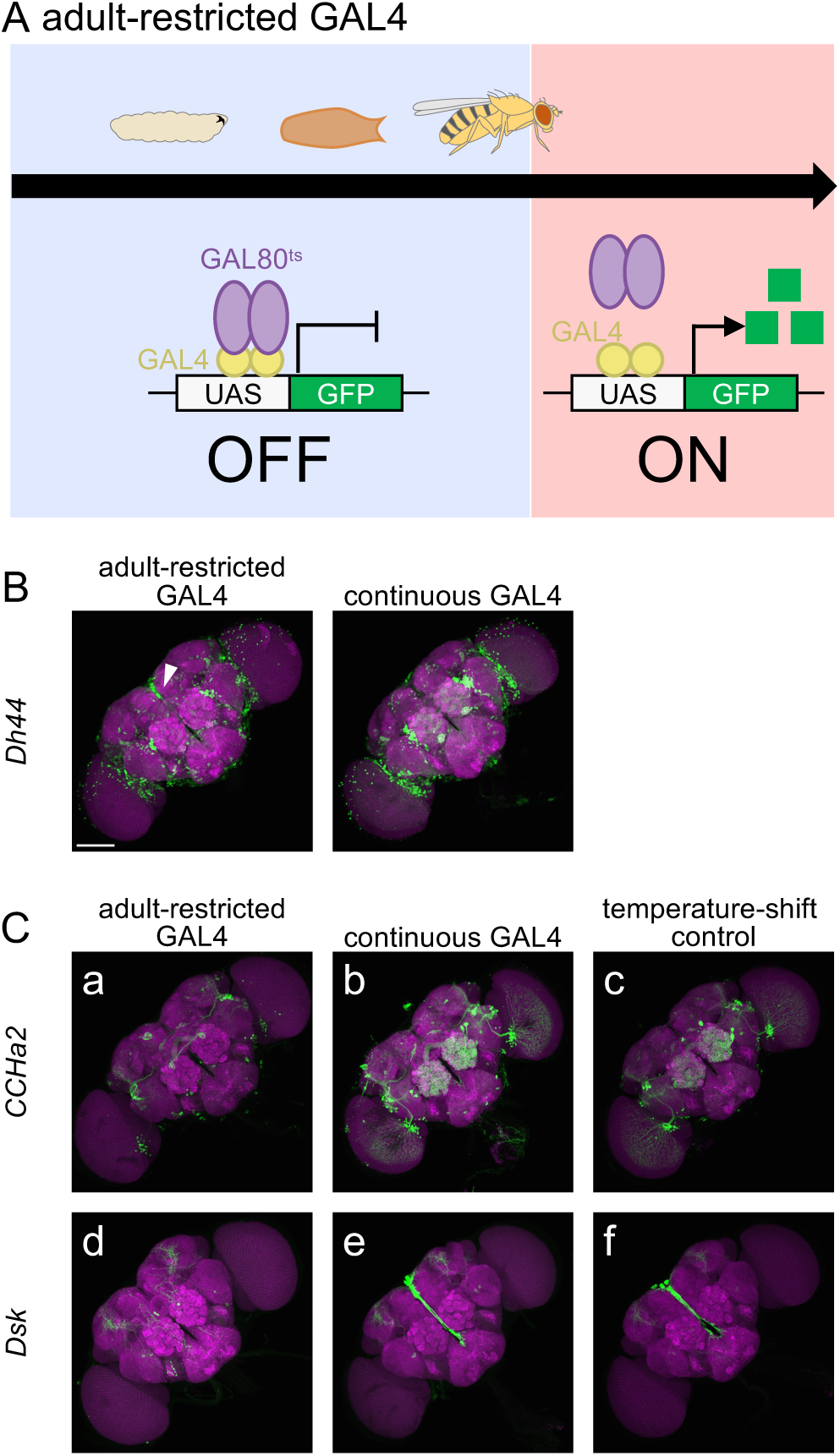
Adult-restricted GAL4 induction reveals condition-dependent reporter labeling in selected neuropeptide-GAL4 knock-in lines. (A) Schematic of the adult-restricted GAL4 experimental design. GAL4 was suppressed by tubP-GAL80ts at 18°C until 4–6 days after eclosion. Flies were then shifted to 30°C for 2 days before dissection to relieve GAL80-mediated suppression. (B) Representative adult brain images of *Dh44-T2A-GAL4* under adult-restricted and continuous GAL4 conditions. The arrowhead indicates additional reporter-positive cells in the pars intercerebralis with ventrally projecting axons that were observed under the adult-restricted condition but not under continuous GAL4 expression. (C) Representative adult brain images of *CCHa2-T2A-GAL4* (a–c) and *Dsk-T2A-GAL4* (d–f). GFP labeling in the antennal lobe of *CCHa2-T2A-GAL4* and in the pars intercerebralis of *Dsk-T2A-GAL4* was observed under continuous GAL4 expression but not under the adult-restricted condition. (a, d) Adult-restricted GAL4 condition. GAL4 activity was suppressed by tubP-GAL80ts at 18°C during development and induced by shifting flies to 30°C after eclosion, as described in (A). (b, e) Continuous GAL4 condition. Flies were reared at 25°C without temporal restriction of GAL4 activity and dissected 5 days after eclosion. (c, f) Temperature-shift controls lacking tubP-GAL80ts, subjected to the same 18°C-to-30°C temperature regimen as in (A). Scale bars, 100 μm.

Using this approach, we compared GFP reporter patterns between continuous and adult-restricted GAL4 conditions in the adult brain and ventral nerve cord (Figures S6 and S7). If continuous GAL4 activity caused cell loss, restricting GAL4 activity to adulthood would be expected to reveal additional cells that were absent under continuous GAL4. We found that in *Dh44-T2A-GAL4*, adult-restricted GAL4 activity revealed additional cells in the pars intercerebralis with ventrally projecting axons, a population of cells previously shown to express Dh44 (Figure 5B; Cannell et al. 2016). This difference is consistent with reduced cell survival or reporter detectability under continuous GAL4 activity. Unexpectedly, we found that GFP labeling in specific cell populations was absent in two lines, *CCHa2-T2A-GAL4* and *Dsk-T2A-GAL4*, when GAL4 activity was restricted to adulthood (Figures 5 and 6). In *CCHa2-T2A-GAL4*, GFP labeling in the antennal lobe was readily detected under continuous GAL4 activity but was not detected following adult-restricted GAL4 induction (Figure 5C-a,C-b). Similarly, *Dsk-T2A-GAL4* produced GFP labeling in the pars intercerebralis with continuous GAL4, whereas this signal was absent following adult-restricted induction (Figure 5C-d,C-e). These observations indicate that adult GFP reporter patterns in these lines depend strongly on the temporal window of GAL4 activity.

**Figure 6.**
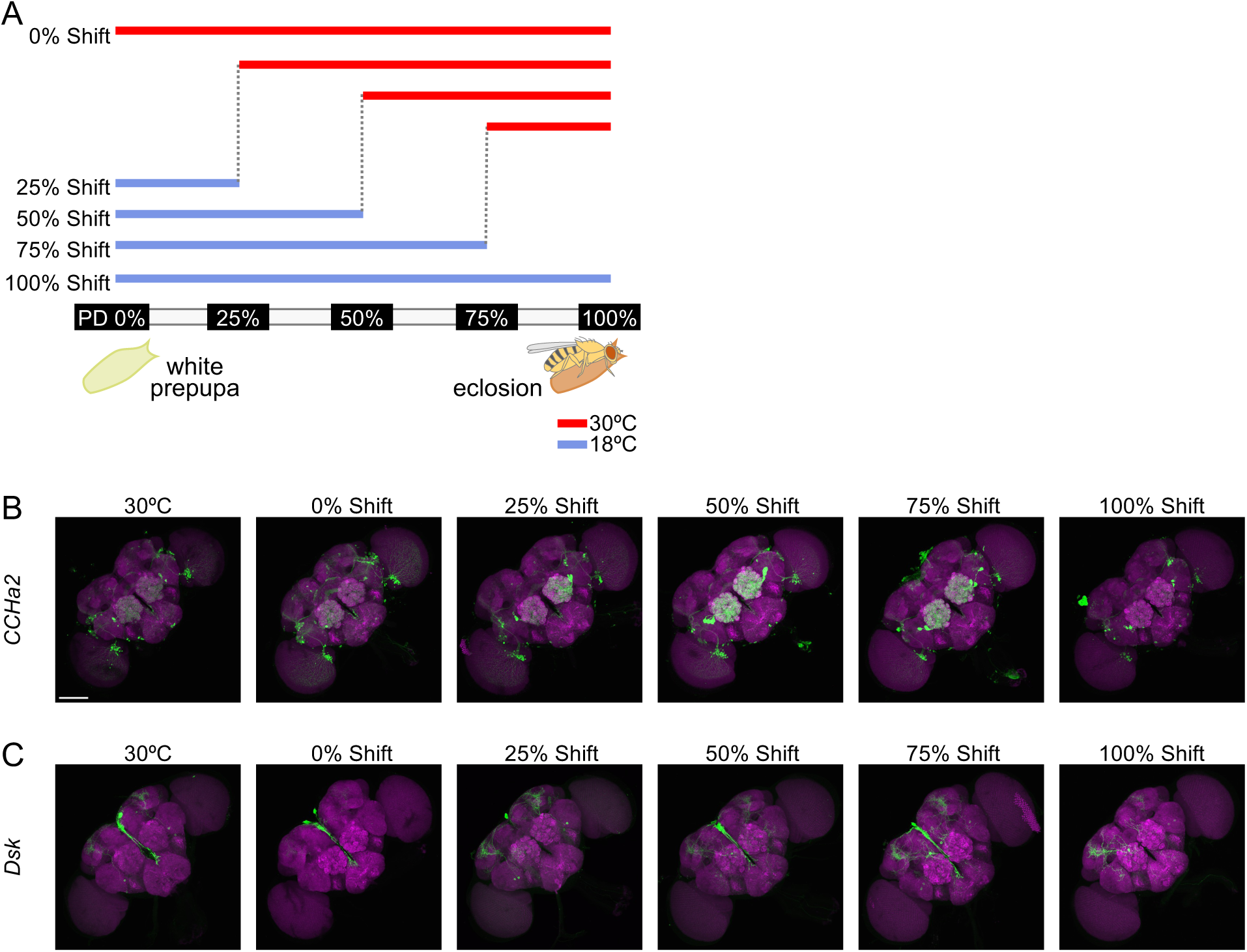
Temporal analysis of GAL4-dependent reporter labeling during pupal development. (A) Schematic of the pupal-stage temperature-shift experiment. White prepupa and the end of pupal development immediately before eclosion were defined as 0% and 100% of pupal development, respectively. At 18°C, development from white prepupa to eclosion took approximately 8 days. For the temperature-shift conditions, flies carrying *tubP-GAL80ts* were shifted from 18°C to 30°C at 48-h intervals, corresponding to approximately 0%, 25%, 50%, 75%, or 100% of pupal development. The 30°C control flies, which did not carry *tubP-GAL80ts*, were reared continuously at 30°C until dissection. (B) Representative adult brain images of *CCHa2-T2A-GAL4* following temperature shifts at the indicated stages of pupal development. (C) Representative adult brain images of *Dsk-T2A-GAL4* following the same temperature-shift regimen. Brains were dissected from female flies 5 days after eclosion. Scale bars, 100 μm.

### 2.5 Pre-eclosion GAL4 activity shapes reporter patterns in the adult brain

The loss of GFP labeling in *CCHa2-* and *Dsk-T2A-GAL4* after adult-restricted GAL4 induction suggested that these adult reporter patterns might originate from GAL4 activity before eclosion. We therefore examined the temporal origin and persistence of these signals in greater detail.

In *CCHa2-T2A-GAL4*, prominent GFP labeling was observed in the antennal lobe under continuous GAL4 activity, consistent with the previously reported innervation of the antennal lobe by CCHa2-expressing neurons (Yamagata et al. 2022), but was absent when GAL4 activity was restricted to adulthood (Figure 5C-a,C-b). To exclude the possibility that this difference was caused by the temperature-shift regimen itself, we subjected animals lacking *tubP-GAL80ts* to the same 18°C-to-30°C shift. Antennal-lobe labeling was retained under this condition (Figure 5C-c), indicating that its loss in the adult-restricted condition resulted from developmental suppression of GAL4 rather than from the temperature shift.

We next asked when the antennal-lobe signal was generated. Using *tubP-GAL80ts*, we shifted animals from 18°C to 30°C at different stages of pupal development and dissected the brains 5 days after eclosion (Figure 6A). Strong antennal-lobe labeling was observed when GAL4 was activated by the 75% point of pupal development, whereas little or no labeling was detected when GAL4 was activated just before eclosion (Figure 6B). These results indicate that the adult antennal-lobe pattern depends on GAL4 activity during pupal development and that GAL4 activity in these cells declines sharply around eclosion.

A similar temporal mismatch was observed with *Dsk-T2A-GAL4*. Under continuous GAL4 activity, GFP-positive cells were consistently detected in the pars intercerebralis, whereas these cells were not labeled when GAL4 activity was restricted to adulthood (Figure 5C-d,C-e). The same temperature-shift regimen in the absence of *tubP-GAL80ts* did not eliminate pars intercerebralis labeling (Figure 5C-f), again excluding temperature treatment itself as the cause of the difference. Pupal-stage induction experiments further showed that this adult reporter pattern depended on GAL4 activity before eclosion (Figure 6A,C). As with *CCHa2-T2A-GAL4*, therefore, the adult *Dsk-T2A-GAL4* reporter pattern retains a substantial contribution from developmental GAL4 activity.

The *CCHa2-* and *Dsk-T2A-GAL4* results demonstrate that stereotyped GFP patterns observed in the adult brain can originate from transient GAL4 activity before eclosion rather than from ongoing adult GAL4 activity.

## 3 Discussion

The neuropeptide T2A-GAL4 lines generated in this study expand the genetic tools available for accessing peptidergic cell populations in *Drosophila*. A variety of neuropeptide-related GAL4 drivers have been developed using promoter fragments, enhancer traps, and knock-ins (Park et al. 2008; Deng et al. 2019), but the availability of these lines from public stock centers remains uneven across neuropeptide genes. Our endogenous T2A-GAL4 knock-ins offer the additional advantage that GAL4 is transcribed from the regulatory environment of the targeted locus, while preserving the coding sequence of the neuropeptide precursor. The lines described here should therefore complement existing neuropeptide driver collections and provide convenient access to peptidergic neurons for anatomical and functional studies. At the same time, our characterization of these lines revealed several practical limitations that are important when GAL4-dependent reporter patterns are used to infer the identity, morphology, or current expression state of labeled cells.

A second useful resource from this study is the collection of fluorescent Brp (Wagh et al. 2006) knock-in alleles (Figure 1B–G). Neuropil counterstaining with the nc82 antibody is widely used to provide anatomical landmarks in the *Drosophila* nervous system, but immunostaining substantially increases sample-processing time. Chemical-tagging approaches such as endogenous Brp-SNAP greatly accelerate this process by replacing antibodies with rapidly diffusing fluorescent ligands (Kohl et al. 2014), but they still require post-fixation labeling and washing steps. By contrast, our fluorescent Brp knock-ins can be imaged directly by native fluorescence without any additional staining or labeling procedure. In particular, Brp-miRFP680 can be combined with GFP- and RFP-based reporters to visualize neuropil landmarks, neuronal membranes and processes. This configuration allowed rapid screening of multiple neuropeptide-GAL4 lines under different experimental conditions. Chemical tags such as SNAP and CLIP retain the advantage that the fluorophore can be selected experimentally and exchanged as imaging requirements change, whereas genetically encoded fluorescent proteins provide a fixed spectral channel (Kohl et al. 2014; Sutcliffe et al. 2017). Thus, the two approaches offer complementary trade-offs between spectral flexibility and experimental simplicity.

Our characterization of the GAL4 lines also reinforces the importance of considering the biological activity of GAL4 itself. Toxic effects of strong GAL4 expression have been recognized for many years and have often been discussed in the context of apoptosis in tissues including imaginal discs and neurons (Abidi and Smith-Bolton 2018; Kramer and Staveley 2003; Rezával et al. 2007). Although we observed increased cell death in tissues expressing strong GAL4 drivers, our results indicate that apoptosis alone does not explain the resulting phenotypes (Figures 3 and 4). The clearest example in this study is *GMR-GAL4*, a classical model of GAL4-associated toxicity. At elevated temperature, *GMR-GAL4* induced substantial cell death in the larval eye disc as well as a rough-eye phenotype in adult flies. Although p35 almost completely suppressed cell death in the eye disc, it failed to restore normal adult eye morphology (Figure 4B,C). Similarly, the developmental phenotypes caused by *Burs/Pburs-T2A-GAL4* and *nub-T2A-GAL4* were poorly rescued by p35 despite being fully suppressible by GAL80 (Figures 3A,B and 4A). Notably, in *Burs-T2A-GAL4*, severe wing expansion defects were not accompanied by a reduction in the number of reporter-labeled Bursicon neurons, although their cell bodies and arbors frequently appeared abnormal. Strong GAL4 activity can therefore induce cell death but can also perturb cellular morphology, differentiation, or physiology without eliminating the affected cells (Figure 3C,D). Consistent with this view, a recent study showed that strong *yolk-GAL4* expression disrupts adult fat-body morphology and multiple physiological functions, whereas inhibition of apoptosis by p35 fails to rescue the physiological defects (Keith et al. 2026).

The molecular basis of these adverse effects of GAL4 remains unclear. At high concentrations, GAL4 might bind cryptic or low-affinity binding sites throughout the *Drosophila* genome and activate the transcription of nearby endogenous genes. Alternatively, excessive GAL4 could sequester limiting components of the transcriptional machinery or transcriptional cofactors, resulting in transcriptional squelching (Gill and Ptashne 1988). These mechanisms are not mutually exclusive and could account for the diverse, tissue-dependent effects of strong GAL4 activity observed here and in previous studies. A different limitation became apparent from the temporal-control experiments. In both *CCHa2-T2A-GAL4* and *Dsk-T2A-GAL4*, specific GFP patterns visible in the adult brain under continuous GAL4 conditions were not reproduced when GAL4 activity was restricted to adulthood (Figure 5C). Pupal-stage induction experiments showed that these patterns depended on GAL4 activity before eclosion (Figure 6). Thus, reporter signals generated during development can remain detectable for at least 5 days after eclosion and contribute substantially to the apparent adult reporter pattern. GFP perdurance has long been recognized as a potential complication in GAL4/UAS-based expression analysis, with developmental studies noting persistent reporter labeling from earlier stages (Soustelle and Giangrande 2007; Manning et al. 2012). More recently, temporal restriction of GAL4 activity revealed substantial age-dependent changes in the activity of several commonly used neuronal and glial drivers (Delandre et al. 2025). A related phenomenon has also been demonstrated for GAL4-driven RNA interference: transient expression of UAS-shRNAs can produce gene knockdown that persists for several days after GAL4 activity has ceased, generating so-called “shadow RNAi” cells (Bosch et al. 2016).

The distinction between ongoing and historical GAL4 activity motivated the development of lineage-tracing systems such as G-TRACE, in which current GAL4 expression is visualized separately from a permanent genetic record of prior GAL4 activity (Evans et al. 2009). Our results show that even a conventional UAS fluorescent reporter, without any dedicated lineage-tracing mechanism, can itself retain enough developmental history to generate a stereotyped adult brain pattern that is no longer reproduced by adult-restricted GAL4 activity. This effect may be particularly pronounced in post-mitotic neurons, where reporter molecules are not diluted by cell division. Thus, in neuronal tissues, the *UAS-CD8::GFP* reporter can unintentionally provide a partial record of earlier driver activity rather than a simple snapshot of ongoing GAL4 expression.

These observations highlight two important considerations for using the GAL4/UAS system, especially in adult post-mitotic neurons. Reporter patterns could reflect GAL4 activity earlier in development, while strong GAL4 activity might alter the number or morphology of labeled neurons. As we have shown, these effects can be readily assessed by temporally restricting GAL4 activity. Such temporal control should therefore improve the accuracy with which GAL4 reporter patterns are interpreted in the adult nervous system.

## 4 Experimental Procedures

### 4.1 Drosophila stocks and husbandry

The following strains were obtained from the Bloomington Drosophila Stock Center (BDSC): *10XUAS-IVS-mCD8::GFP* (#32185), *UAS-histone-RFP* (#56555), R13F02-GAL4 (#48571). The following strains were obtained from the Kyoto Stock Center (DGRC): *UAS-p35* (#108-018), *tubP-GAL80ts* (#130-455). The third-chromosome *GMR-GAL4^J14^* (Freeman 1996) line was obtained from the laboratory of Dr. Yasushi Hiromi.

Flies were maintained on standard cornmeal-based food at 25°C unless otherwise indicated. For experiments using temperature-sensitive GAL80, flies were reared at 18°C and shifted to 30°C to induce GAL4 activity.

### 4.2 Generation of transgenic flies

Fluorescent Brp knock-in lines were generated from the previously established *brp-GFP11* allele (Wu et al. 2025) by phiC31-mediated cassette exchange. Exchange vectors were constructed by cloning the coding sequences of *Venus*, *mBaoJin*, *mScarlet-I*, *mTurquoise2*, *miRFP680*, or *mGarnet2* into *pF3BGX* (Kondo et al. 2020). Each exchange plasmid was injected into fertilized eggs carrying the *brp-GFP11* allele and *nos-phiC31* (BDSC #34770) to mediate attP/attB recombination at the *brp* locus. Transformants were visually identified by the eye-specific *3xP3-Venus* fluorescent marker carried by the exchange cassette. The parental *brp-GFP11* allele also contained a floxed *3xP3-RFP* marker at the *brp* locus. Unless otherwise indicated, both fluorescent markers were subsequently removed by crossing to the *CyO hs-Cre* balancer (BDSC #1092). In the Brp-Venus and Brp-mBaoJin lines shown in Figure 1E,F, both markers were deliberately retained to demonstrate simultaneous imaging of the Brp fusion proteins and RFP. The resulting alleles encode fluorescent proteins fused to the C-terminus of endogenous Brp.

Neuropeptide T2A-GAL4 knock-in lines and *nub-T2A-GAL4* were generated essentially as previously described (Kondo et al. 2020). The sequences of primers used to amplify the homology arms for each donor vector and oligonucleotides used to construct the gRNA expression vectors are listed in Table S1. Donor and corresponding gRNA plasmids were co-injected into fertilized eggs of the *nos-Cas9* strain (Kondo and Ueda 2013). Transformants were identified by the eye-specific *3xP3-RFP* fluorescent marker and established as balanced stocks. Unless otherwise indicated, the *3xP3-RFP* marker was subsequently removed by crossing to *hs-Cre* balancers (BDSC #1092 and #1501).

The *tubP-GAL80* line used in Figures 3, 4 and S5 was generated by phiC31-mediated integration of a *tubP-GAL80* construct into the attP40 landing site (BDSC #79604). The *tubP-GAL80* construct contained the *αTub84B* promoter, the GAL80 coding sequence, and the p10 polyadenylation signal.

### 4.3 Fly crosses and temperature-shift experiments

For systematic expression profiling of the neuropeptide T2A-GAL4 knock-in lines, each GAL4 line was crossed to a reporter stock carrying *brp-miRFP680*, *UAS-CD8::GFP*, and *UAS-histone-RFP*. The resulting progeny, referred to as the continuous GAL4 condition, were reared at 25°C throughout development and after eclosion and dissected 5 days after eclosion. For adult-restricted GAL4 experiments, each GAL4 line was crossed to a reporter stock carrying *brp-miRFP680*, *tubP-GAL80ts*, *UAS-CD8::GFP*, and *UAS-histone-RFP*. These flies were reared at 18°C throughout development, maintained at 18°C for 4–6 days after eclosion, and then shifted to 30°C for 2 days before dissection. Temperature-shift controls lacking *tubP-GAL80ts* were subjected to the same temperature regimen as the adult-restricted GAL4 condition.

To examine GAL4-associated phenotypes, *Burs-T2A-GAL4*, *Pburs-T2A-GAL4*, *nub-T2A-GAL4*, and *GMR-GAL4* lines were crossed to *yw* control flies or to flies carrying *tubP-GAL80* or *UAS-p35*, as indicated. For experiments requiring selection of GAL4-carrying larvae, the GAL4 chromosome was maintained over either *TM6B, Tb^1^* (BDSC #2995) or *CyO, P{2xTb^1^-RFP}* (BDSC #36336), allowing GAL4-carrying larvae to be identified by the absence of the *Tubby* phenotype following experimental crosses. For pupal-stage temperature-shift experiments, the duration of pupal development was normalized from 0% at the white prepupal stage to 100% immediately before eclosion. Animals were shifted from 18°C to 30°C at the indicated percentages of pupal development and maintained at 30°C until dissection.

### 4.4 Dissection, immunohistochemistry, and fluorescence imaging

Adult brains and VNCs were dissected from virgin females, and late-pupal VNCs were dissected from female pupae. Larval CNSs were also dissected, and all dissections were performed in phosphate-buffered saline (PBS). Larval CNSs were fixed in 4% paraformaldehyde (PFA) in PBS for 15 min at room temperature (RT), whereas adult brains, adult VNCs, and late-pupal VNCs were fixed in the same fixative for 40 min at RT. Following fixation, samples were washed three times for 10 min each in 0.1% Triton X-100 in PBS (PBST) at RT. Larval preparations were stored and mounted in 80% glycerol. Adult brain, adult VNC, and late-pupal VNC preparations were stored in 25% glycerol and subsequently mounted in Omnipaque (GE Healthcare) for confocal imaging.

For immunostaining of wing imaginal discs, discs were dissected and fixed in 4% PFA for 15 min at RT with gentle agitation. After fixation, the discs were washed three times in PBST and incubated in 4% normal goat serum (NGS) (Sigma-Aldrich, #G9023-10ML) in PBST for 30 min at RT. Samples were then incubated overnight at 4°C with primary antibodies diluted in 4% NGS/PBST. The primary antibodies used were mouse anti-Wg (1:150; DSHB, #4D4) and rabbit anti-cleaved Dcp-1 (1:500; Cell Signaling Technology, #9578). The following day, samples were washed three times for 10 min each in PBST at RT and incubated with secondary antibodies diluted in 4% NGS/PBST for 2 h at RT. Alexa Fluor 647 donkey anti-mouse IgG (1:700; Jackson ImmunoResearch, #715-605-151) and Cy3 donkey anti-rabbit IgG (1:1,000; Jackson ImmunoResearch, #711-165-152) were used as secondary antibodies. Samples were then washed three times for 10 min each in PBST, mounted in 80% glycerol, and stored at 4°C until confocal imaging.

Native fluorescence from CD8::GFP, histone-RFP and fluorescent Brp knock-ins was imaged using an FV3000 confocal microscope (Olympus/Evident). Maximum-intensity projections were generated from confocal z-stacks for the representative images shown. Acquisition settings were optimized for each experiment and were not necessarily identical across experiments. For quantitative comparisons of fluorescence intensity or photobleaching, all samples within each experiment were imaged using identical acquisition settings.

### 4.5 Adult wing and eye imaging

Adult wings were dissected, rinsed in 100% ethanol, mounted in silicone oil (Shin-Etsu Chemical, #KF-96-20CS), and imaged using an Axio Imager A2 microscope (Carl Zeiss). Adult eyes were imaged using an SZX16 stereomicroscope (Olympus).

### 4.6 Acridine Orange staining

Eye imaginal discs were dissected from larvae of the indicated genotypes and incubated in 0.5 μg/mL Acridine Orange (Tokyo Chemical Industry, #A3396) diluted in PBS for 15 min at RT. Following staining, the discs were briefly rinsed in PBS, mounted in PBS and imaged using a ZOE Fluorescent Cell Imager (Bio-Rad).

### 4.7 Photobleaching assay

Photobleaching was assessed by repeatedly imaging the same region for 1,000 consecutive scans under identical confocal acquisition settings. Fluorescence intensity within the indicated region of interest (Figure S2D) was measured throughout the image series and normalized to the initial fluorescence intensity.

### 4.8 Statistical analysis

Statistical significance was assessed using two-sided Welch’s *t*-tests. Statistical analyses were performed and graphs were generated using BioChart v0.1.1, custom-written JavaScript software (https://github.com/tus-kondolab/biochart).

## Author contributions

**Kyoko Jinnai**: conceptualization, investigation, formal analysis, writing – original draft, writing – review and editing. **Kota Ozawa**: investigation. **Katsuki Yajima**: investigation. **Hiromu Tanimoto**: conceptualization, funding acquisition. **Shu Kondo**: conceptualization, methodology, investigation, supervision, project administration, funding acquisition, writing – original draft, writing – review and editing.

## Acknowledgments

We thank the Bloomington *Drosophila* Stock Center, the Kyoto *Drosophila* Stock Center and the NIG-Fly Stock Center for providing fly stocks. ChatGPT (OpenAI; GPT-5.5, GPT-5.6 and GPT-6 Pro) was used to refine the wording and improve clarity in the Abstract, Introduction, Results, Discussion, Experimental Procedures and figure legends.

## Ethics Statement

Not applicable.

## Data Availability Statement

The neuropeptide T2A-GAL4 and fluorescent *brp* knock-in lines will be deposited at the Kyoto Stock Center upon acceptance of the manuscript. The data supporting the findings of this study and other reagents generated in this study, including plasmids, are available from the corresponding author upon reasonable request. BioChart, the custom software used for statistical analysis and graph generation, is publicly available at https://github.com/tus-kondolab/biochart.

## Final Approval

All authors have read and approved the final manuscript and agree to its submission.

