## Supplementary Figures for "Neuropeptide T2A-GAL4 knock-ins illustrate practical considerations for interpreting GAL4 reporter patterns in *Drosophila*"

### Figure S1

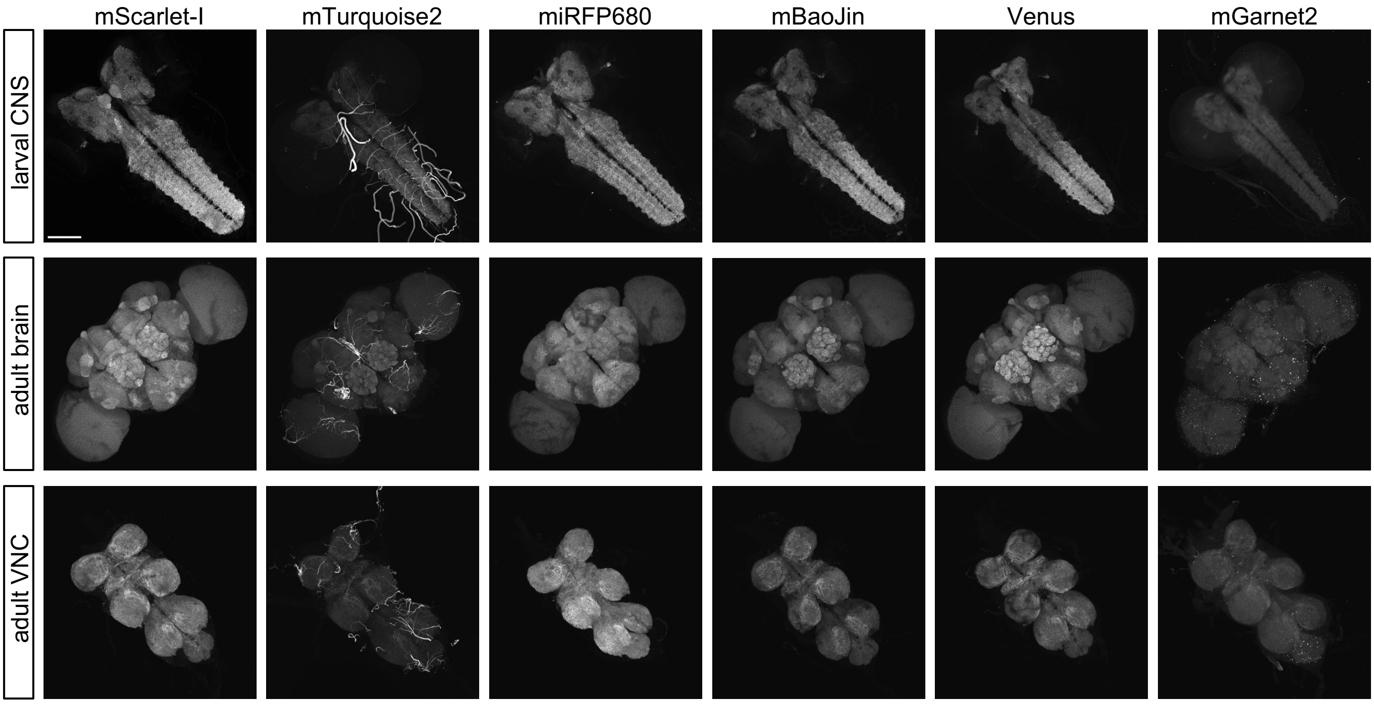


**Figure S1. Fluorescent Brp labeling in the larval CNS, adult brain, and adult VNC.** Representative images of Brp-Venus, Brp-mBaoJin, Brp-mScarlet-I, Brp-mTurquoise2, Brp-miRFP680, and Brp-mGarnet2 in CNSs from third-instar larvae and in brains and VNCs from 3-day-old adult females. Scale bars, 100 μm.

### Figure S2

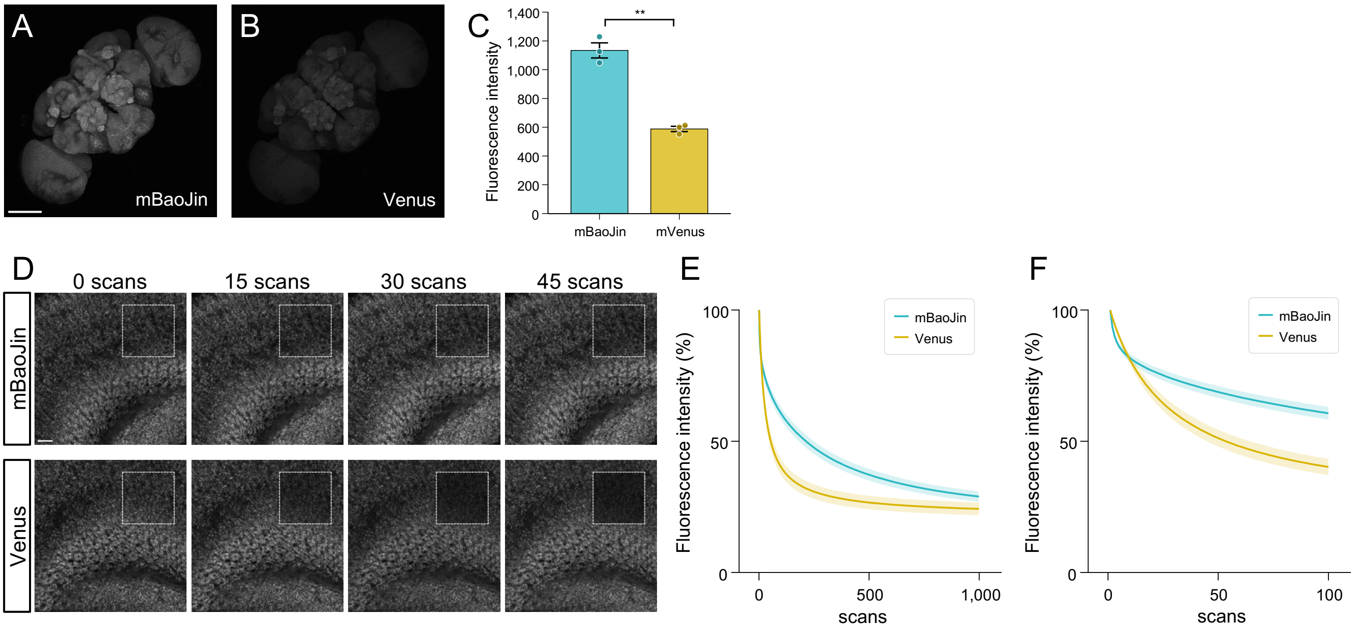


**Figure S2. Brp-mBaoJin provides a bright and photostable neuropil reference signal.**

(A,B) Representative adult female brain images of Brp-Venus and Brp-mBaoJin acquired under identical imaging conditions. Scale bars, 100 µm. (C) Quantification of fluorescence intensity measured from defined brain ROIs in Brp-Venus and Brp-mBaoJin flies. Data are presented as mean ± SEM with individual data points; n = 3 brains per genotype. Statistical significance was assessed using Welch's t-test. **p < 0.01. (D) Single confocal images showing photobleaching of Brp-Venus and Brp-mBaoJin during repeated scanning. Images acquired after 0, 15, 30 and 45 scans are shown. Scale bars, 10 µm. (E–F) Quantification of photobleaching during repeated scanning. Fluorescence intensity was measured from the same defined ROI and normalized to the initial intensity. (E) Fluorescence intensity over 1,000 scans. (F) Fluorescence intensity over the first 100 scans. Solid lines indicate the mean, and shaded areas indicate ± SEM (n = 3 brains per genotype).

Figure S3

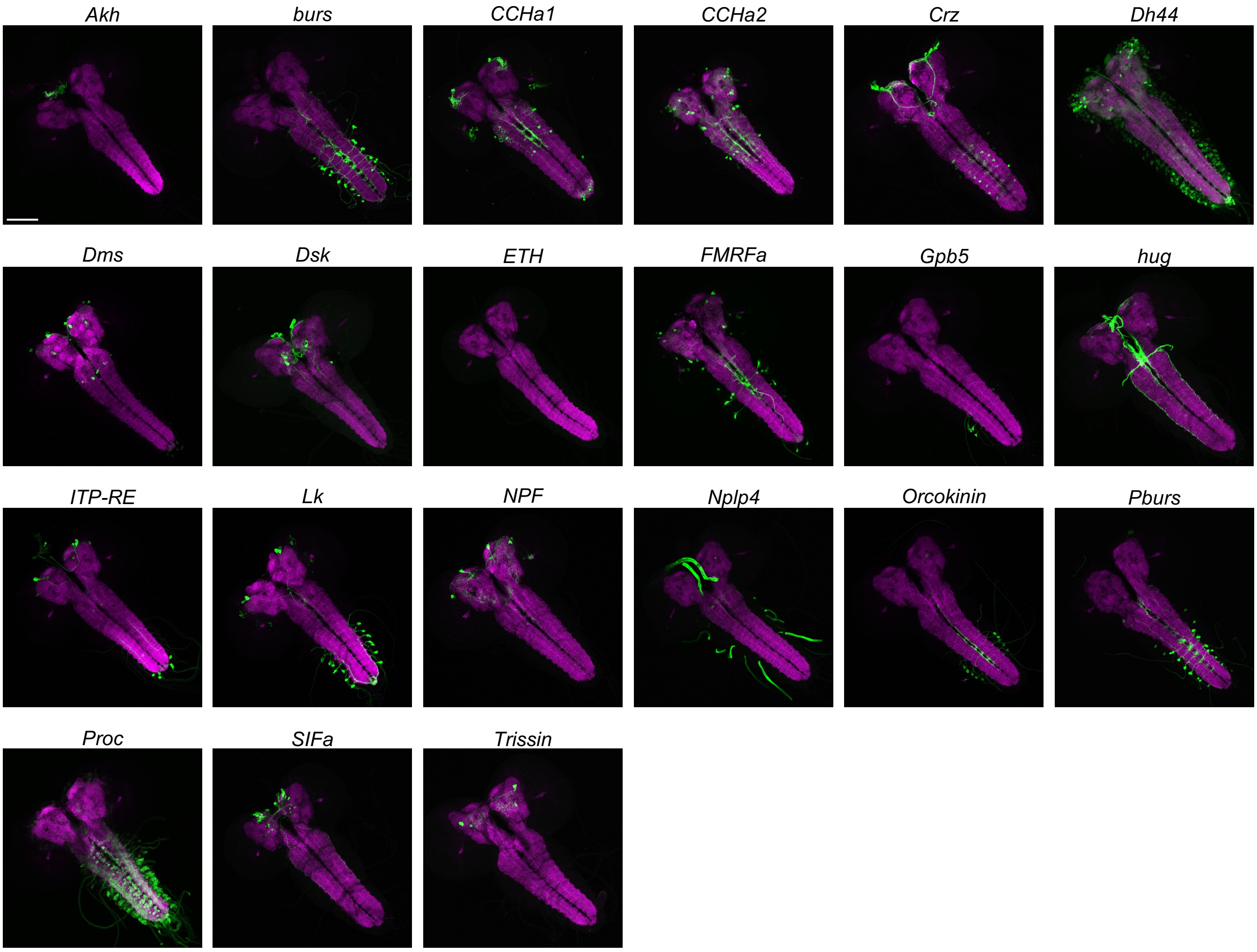


**Figure S3. Larval CNS reporter labeling patterns of neuropeptide-GAL4 knock-in lines.**

Representative larval CNS montage of the indicated neuropeptide-GAL4 lines. CD8::GFP (green) provides GAL4-dependent membrane labeling, while Brp-miRFP680 (magenta) provides neuropil landmarks. Larval CNSs were obtained from third-instar larvae. Scale bar, 100 μm.

### Figure S4

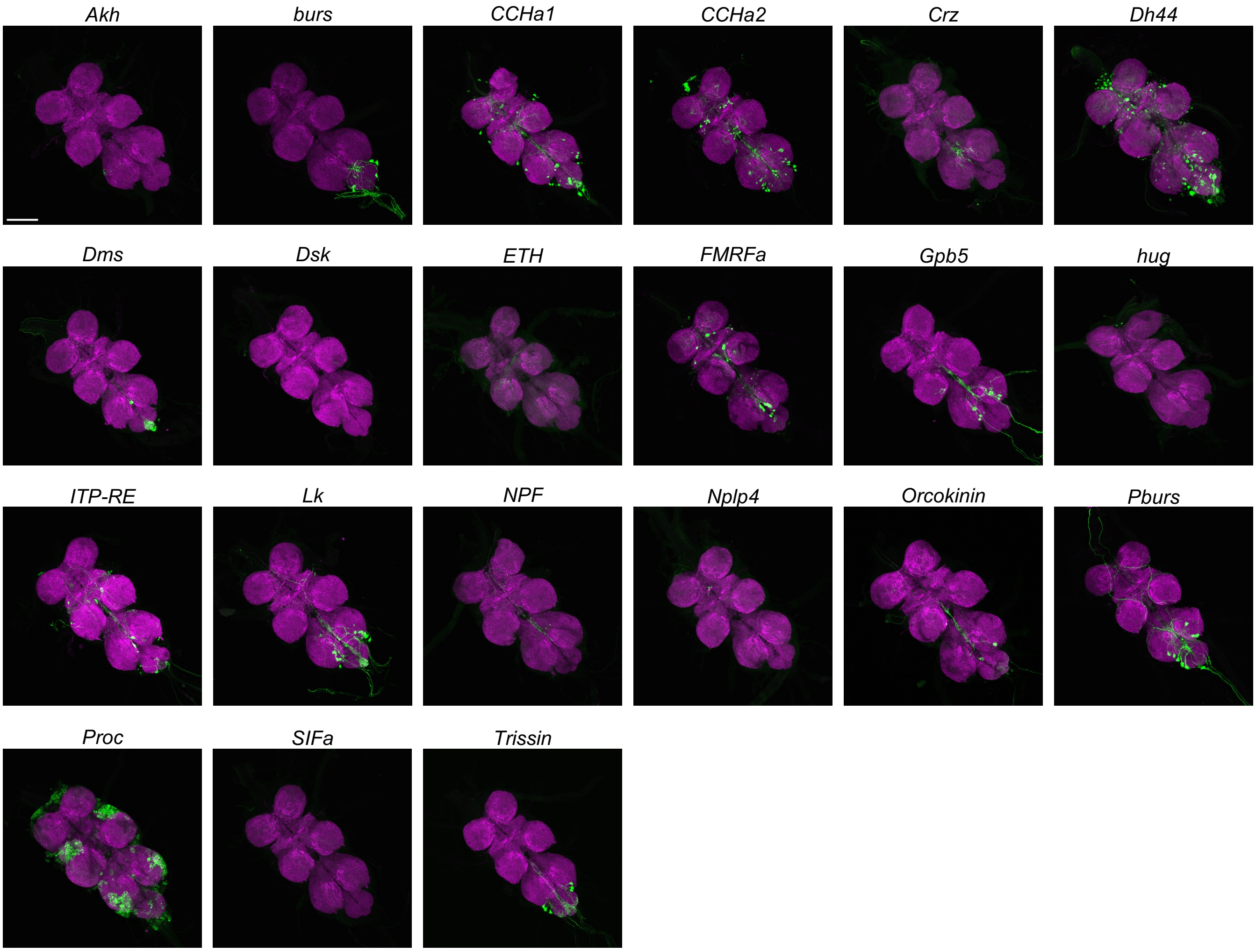


**Figure S4. Adult VNC reporter labeling patterns of neuropeptide-GAL4 knock-in lines.**
Representative adult VNC montage of the indicated neuropeptide-GAL4 lines. CD8::GFP (green) provides GAL4-dependent membrane labeling, while Brp-miRFP680 (magenta) provides neuropil landmarks. Adult VNCs were obtained from 5-day-old female flies. Scale bar, 100 µm.

### Figure S5

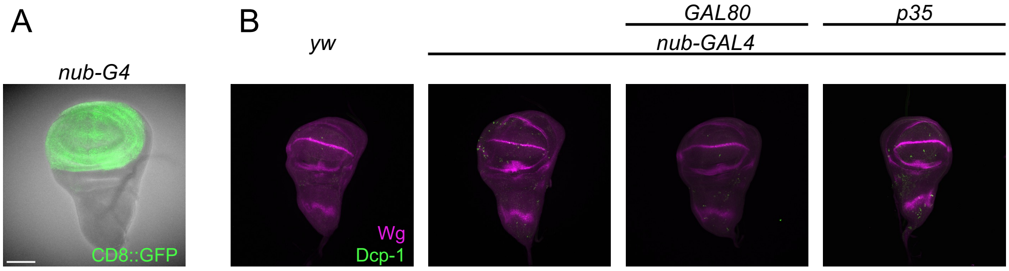


**Figure S5. Effects of *nub-T2A-GAL4* on wing imaginal discs.**
(A) Representative third-instar larval wing imaginal disc showing the expression pattern of *nub-T2A-GAL4* visualized with *UAS-CD8::GFP* (green). (B) Representative third-instar larval wing imaginal discs from *yw*, *nub-T2A-GAL4*, *nub-T2A-GAL4/tubP-GAL80*, and *nub-T2A-GAL4/UAS-p35*, immunostained for cleaved Dcp-1 (green) and Wingless (Wg; magenta). Larvae were reared at 25°C. Scale bar, 100 µm.

### Figure S6

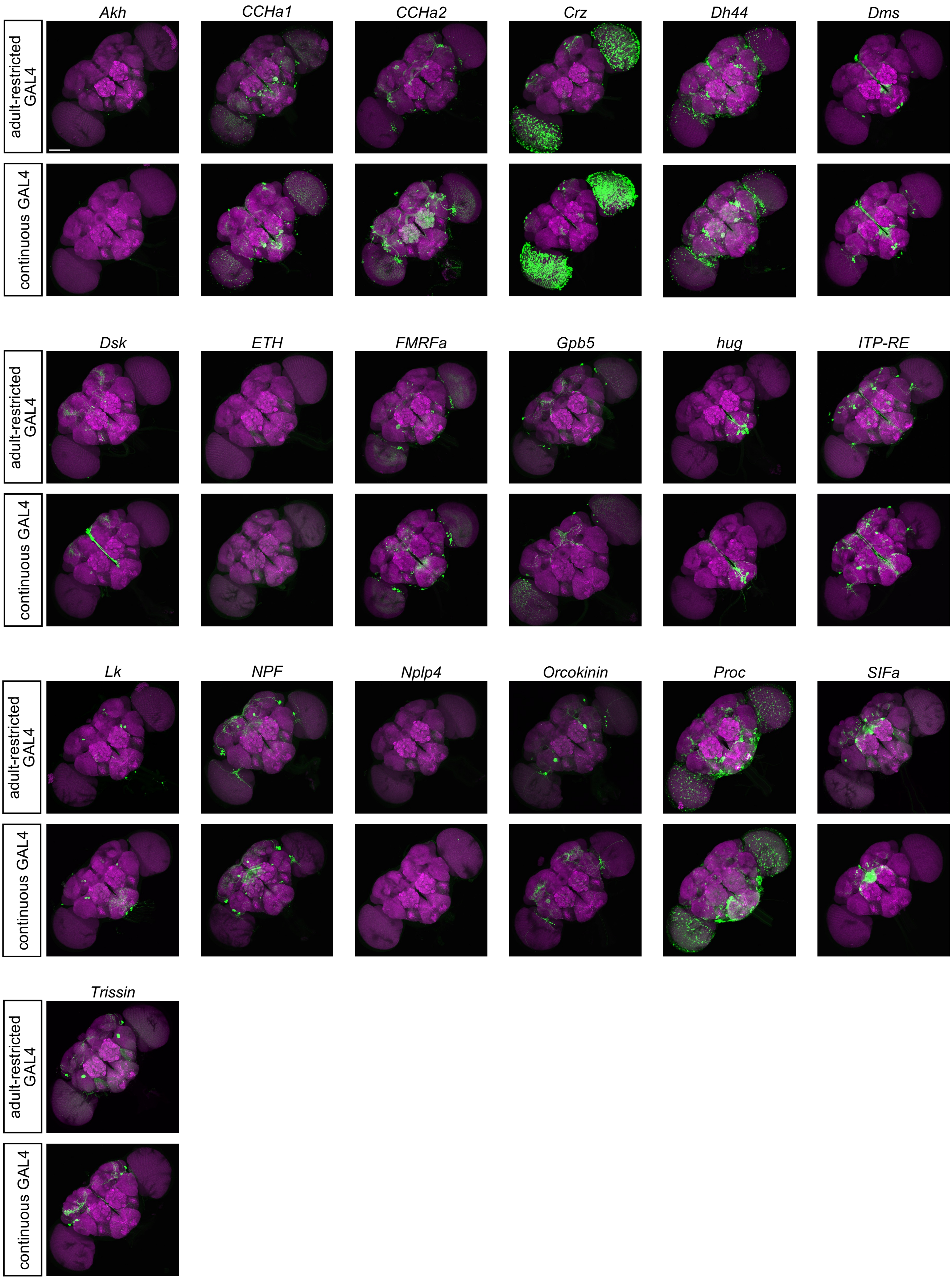
Akh
CCHa1
CCHa2
Crz
Dh44
Dms
adult-restricted GAL4
continuous GAL4
Dsk
ETH
FMRFa
Gpb5
hug
ITP-RE
Lk
NPF
Nplp4
Orcokinin
Proc
SIFa
Trissin

**Figure S6. Comparison of adult-restricted and continuous GAL4 reporter labeling in the brain of neuropeptide-GAL4 lines.**

Representative adult brain images of the indicated neuropeptide-GAL4 lines under adult-restricted GAL4 and continuous GAL4 conditions, shown side by side for comparison. CD8::GFP (green) provides GAL4-dependent membrane labeling, while Brp-miRFP680 (magenta) provides neuropil landmarks. Adult-restricted GAL4 samples were dissected after the 18°C-to-30°C temperature-shift regimen described in Figure 5A, whereas continuous GAL4 samples were maintained at 25°C and dissected 5 days after eclosion. Scale bar, 100 μm.

### Figure S7

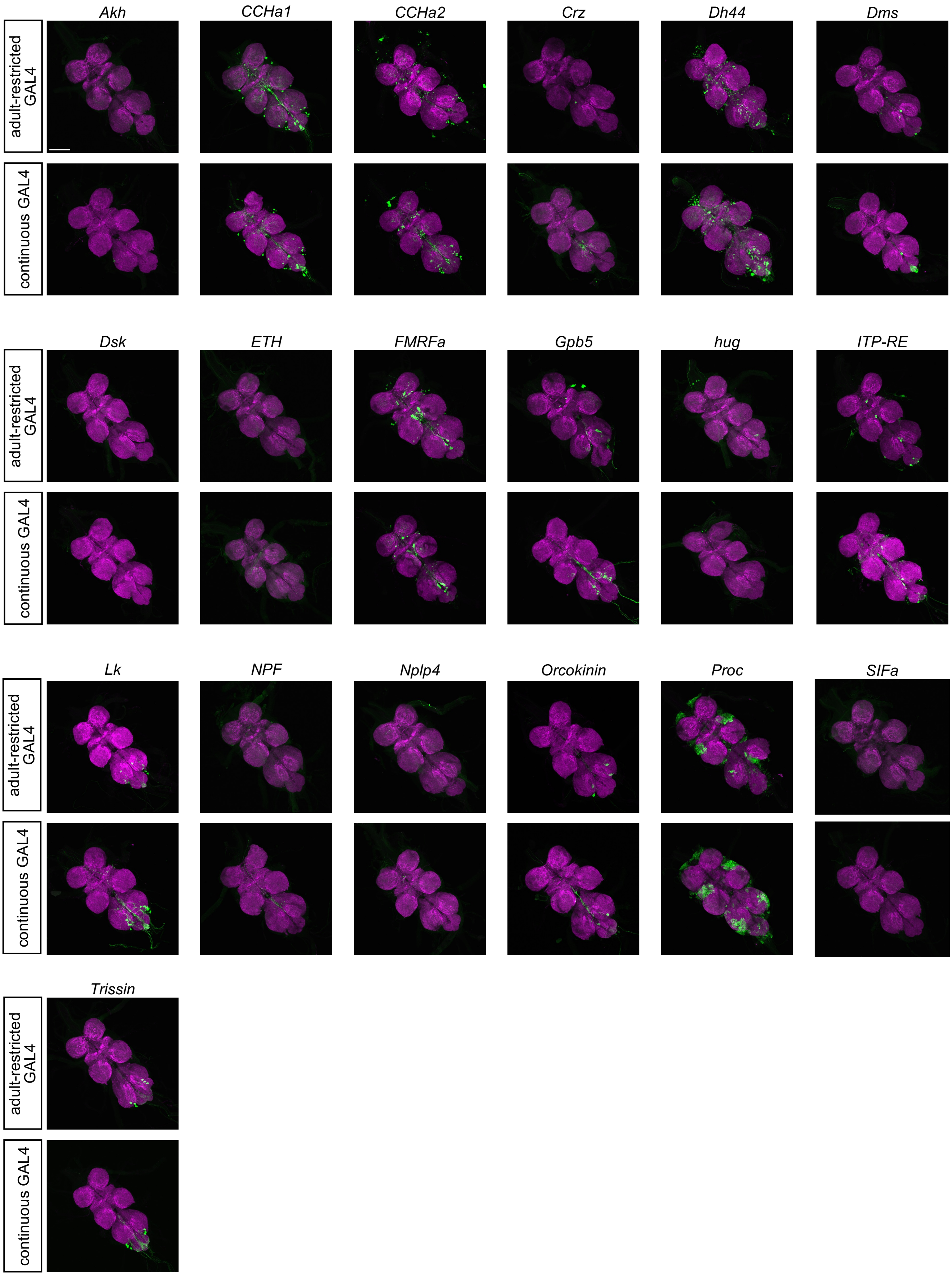
Akh
CCHa1
CCHa2
Crz
Dh44
Dms
adult-restricted GAL4
continuous GAL4
Dsk
ETH
FMRFa
Gpb5
hug
ITP-RE
Lk
NPF
Nplp4
Orcokinin
Proc
SIFa
Trissin

**Figure S7. Comparison of adult-restricted and continuous GAL4 reporter labeling in the VNC of neuropeptide-GAL4 lines.**

Representative adult VNC images of the indicated neuropeptide-GAL4 lines under adult-restricted GAL4 and continuous GAL4 conditions, shown side by side for comparison. CD8::GFP (green) provides GAL4-dependent membrane labeling, while Brp-miRFP680 (magenta) provides neuropil landmarks. Adult-restricted GAL4 samples were dissected after the 18°C-to-30°C temperature-shift regimen described in Figure 5A, whereas continuous GAL4 samples were maintained at 25°C and dissected 5 days after eclosion. Scale bar, 100 μm.
